# Aspirin inhibition of prostaglandin synthesis impairs egg development across mosquito taxa

**DOI:** 10.1101/2020.07.17.208389

**Authors:** Md. Abdullah Al Baki, Shabbir Ahmed, Hyeogsun Kwon, David R. Hall, Ryan C. Smith, Yonggyun Kim

**Affiliations:** Department of Plant Medicals, College of Life Sciences, Andong National University, Andong 36729, Korea; Department of Entomology, Iowa State University, Ames, Iowa 50011, USA

## Abstract

Several endocrine signals are known to mediate mosquito egg development including insulin-like peptide, 20-hydroxyecdysone, and juvenile hormone. The objective of this study was to determine the effects of prostaglandin E_2_ (PGE_2_) as an additional mediator of oogenesis in the mosquitoes, *Aedes albopictus* and *Anopheles gambiae*. The injection of aspirin (an inhibitor of cyclooxygenase) shortly after blood-feeding significantly inhibited egg development at choriogenesis in a dose-dependent manner in *Ae. albopictus*. Moreover, oral administration of aspirin to *An. albopictus* and *An. gambiae* also inhibited egg production. The aspirin treatment suppressed expression of the genes (*Yellow-g* and *Yellow-g2*) associated with exochorion darkening and led to the production of a malformed egg shell in *Ae. albopictus*. These inhibitory effects of aspirin on egg development were rescued by the addition of PGE_2_, confirming the specificity of aspirin in inhibiting prostaglandin production. To validate these results, we identified a putative PGE_2_ receptor (Aa-PGE_2_R) in *Ae. albopictus. Aa-PGE*_*2*_*R* expression was highly inducible in adult ovary after blood-feeding. RNA interference of *Aa-PGE*_*2*_*R* expression resulted in the significant suppression of choriogenesis similar to aspirin treatment, where the addition of PGE_2_ to *Aa-PGE*_*2*_*R*-silenced females failed to rescue egg production. Together, these results suggest that PG synthesis and signaling are required for egg development across diverse mosquito taxa.

**Author Summary:** Progstaglandins (PGs) play crucial roles in mediating various physiological processes in insects. Aspirin (ASP) inhibits PG biosynthesis and has been used as an anti-inflammatory drug. ASP injection or feeding to mosquitoes of *Aedes albopictus* or *Anopheles gambiae* significantly inhibits egg production at chorion formation. This led to significant reduction in fecundity and egg hatchability. PG signal is interrupted by RNA interference (RNAi) of PGE_2_ receptor. The RNAi treatment also gave a similar damage to females in egg production as seen in ASP treatment. Thus, PG signal is required for egg production of these mosquitoes.

**Data Availability Statement:** All relevant data are within the manuscript and its Supporting Information file.

## Introduction

Mosquito-borne diseases are recognized as a leading killer of humans worldwide. The Asian Tiger mosquito, *Aedes albopictus*, transmits at least 22 arboviruses as a capable vector causing Chikungunya, Dengue, and Zika viruses [1,2]. The African Malaria vector mosquito, *Anopheles gambiae*, transmits malaria parasites and results in more than one million annual deaths [3].

Female mosquitoes depend on blood-feeding to initiate egg development mediated by several endocrine signals [4]. Insulin-like peptides can sense nutrient availability and stimulate proliferation of germline stem cells [5]. Juvenile hormone (JH) acts as a main gonadotropic endocrine signal by activating vitellogenesis synthesis from fat body and subsequent vitellogenin (Vg) uptake by growing oocytes [6]. In mosquitoes, 20-ydroxyecdysone (20E) plays a crucial role in stimulating Vg synthesis after blood-feeding [7]. Prostaglandins (PGs) have also been implicated stimulating oviposition in some crickets, in which PG-synthetic machinery is delivered from males during mating to produce massive amounts of PGs in female spermatheca [8]. The reproductive role of PGs is extended to mediate oogenesis in a fruit fly, *Drosophila melanogaster*, in which PGE_2_ facilitates a nurse cell-dumping process to growing oocytes [9]. In addition, PGE_2_ can mediate egg shell protein synthesis during choriogenesis of several insects [10-12].

PGs are a group of oxygenated C20 polyunsaturated fatty acid usually derived from arachidonic acid (AA) [13]. AA is oxygenated by cyclooxygenase (COX) to form various PGs in vertebrates [14]. In insects, specific peroxidases called peroxinectin [15,16] and heme peroxidase [17] are likely to act like COX, with additional specific PG synthases that catalyze the isomerization from PGH_2_ to PGE_2_ or PGD_2_ [18,19]. PGE_2_ plays a role in mediating cellular and humoral immune responses of several mosquitoes [17,20]. However, little is known about the role of PG in mosquito reproduction.

In this study, we examine the effects of PG on *Ae. albopictus* and *An. gambiae* oocyte development using aspirin, a potent anti-inflammatory drug that can inhibit PG biosynthesis [21]. Through the use of aspirin feeding or injection to suppress PG production, we demonstrate a significant reduction in mosquito oocyte development. These phenotypes were further validated by genetically manipulation of PG signaling. The *Ae. albopictus* PGE_2_ receptor was identified and its expression was suppressed by RNA interference (RNAi), enabling experiments to assess PGE_2_ receptor function on oocyte development. The addition of PGE_2_ on RNAi-treated mosquito females was unable to rescue prostaglandin signaling, confirming the specificity of the PGE_2_ receptor.

## Methods

### Insect rearing

*Ae. albopictus* was maintained in laboratory conditions: 27°C temperature, 70 ± 10% relative humidity, and 12:12 h (L:D) photoperiod. First instar larvae were reared in a plastic box (12 8 × 5 cm) containing 0.5∼1.0 L of distilled water with fish food mini pellet (Ilsung, Daegu, Korea). A 10% sucrose solution was supplied to adults using a cotton braid. Female mosquitoes (4 ∼ 5 days old post-adult emergence (PE)) were fed blood from an out-bred mouse line (SLC, Shizuoka, Japan) twice a week to initiate egg development. Each blood-feeding (BF) took 1 h in the afternoon (2 ∼ 6 pm) at photophase. For females to oviposit, a wet kitchen towel was placed on a petri dish (8 cm in diameter) filled with distilled water.

*An. gambiae* (Keele strain) were reared at 27°C and 80% relative humidity, with a 14/10 h day/night cycle. Larvae were fed on fish flakes (Tetra, Blacksburg, VA, USA), while adults were maintained on a 10% sucrose solution.

### Chemicals

Aspirin (ASP: 2-acetoxybenzoic acid), dexamethasone (DEX: (11β,16α)-9-fluoro-11,17,21-trihydroxy-16-methylpregna-1,4-diene-3), ibuprofen (IBU: α-methyl-4-isobutyl phenylacetic acid), prostaglandin E_2_ (PGE_2_: (5Z,11α,13E,15S-11,15-dihydroxy-9-oxoprosta-5,13-dienoic acid), naproxen (NAP: 6-methoxy-α-methyl-2-naphthaleneacetic acid), leukotriene B4 (LTB_4_: 5,12-dihydroxy-6,8,10,14-eicosatetraenoic acid), and 14,15-epoxyeicosatrienoic acid (14,15-EET) were purchased from Sigma-Aldrich Korea (Seoul, Korea) and dissolved in dimethyl sulfoxide (DMSO). For experiments with *An. gambiae*, aspirin was purchased from Cayman Chemical (Ann Arbor, MI, USA).

### Microinjection of test chemicals to mosquito females

For chemical treatment, females (10 min before BF) were anesthetized with CO_2_ gas for 1 min. These females were then injected with 1 μL of test chemical with a glass capillary using a sutter CO_2_ picopump injector (PV830, World Precision Instrument, Sarasota, FL, USA) under a stereomicroscope (SZX-ILLK200, Olympus, Tokyo, Japan). Before injection, micro capillaries (10 µL quartz, World Precision Instrument) with sharp points (< 20 µm in diameter) were prepared with a Narishige magnetic glass microelectrode horizontal puller model PN30 (Tritech Research, Inc., Los Angeles, CA, USA).

### ASP injection and rescue experiment with PGE_2_ or other eicosanoids

One microliter of ASP (1 μg/μL) was injected to each female at 5 days old after adult emergence. For rescue experiment, another 1 μL of PGE_2_, 14,15-EET, or LTB_4_ (10 μg/μL) was injected to each female at 12 h after ASP treatment. Treated females were blood-fed. After 3 days, whole ovaries were dissected in 100 mM phosphate-buffered saline (PBS, pH 7.4) under a stereomicroscope (Stemi SV11, Zeiss, Germany). Each treatment was replicated with 10 females.

### Feeding mosquito females with ASP

Different concentrations of ASP were mixed with 10% sucrose solution and provided to adult female *Ae. albopictus* or *An. gambiae* for sugar-feeding. For *Ae. albopictus* experiments, sucrose (control) or ASP (1 µg/µL) in sucrose solution (5 mL) was placed in a vial with cotton plug and provided to an adult cage containing 10 females 3 h before BF. BF was performed using a mouse in the cage for 1 h. All mice were reared under pathogen-free condition and were anaesthetized using CO_2_ during the blood feeding experiment. The mice used in this study were not euthanized. After BF, treated female mosquitoes were maintained on the sucrose or sucrose-ASP solution for an additional 3 days before ovary dissection. For experiments with *An. gambiae*, newly emerged mosquitoes (0-1 day old) were fed with either aspirin (10 mg/mL) in 10% sucrose solution or 10% sucrose solution (control). After three days female mosquitoes were fed on an anaesthetized female Swiss Webster mouse and continually provided with either sucrose solution before dissection 72 h post BF (PBF).

### Counting oocytes at different developmental stages

For oocyte counting, females were allowed to feed blood for 1 h in the afternoon (2-6 pm) at 3 days PE. Females were mated with males of the same age after BF and reared with 10% sugar solution. After BF, the total number of oocytes was counted up to 4 days. To determine the effect of aspirin on *Ae. albopictus* ovary development, ovary development was assessed up to 4 days PBF under a stereo microscope (Stemi SV11, Zeiss, Germany). Individual ovary follicles were measured under a stereo microscope. For experiments with *An. gambiae*, the total number of developed oocytes were counted with a stereo microscope (Nikon, Melville, NY, USA) and images of the dissected ovaries were taken with a Zeiss SteREO Discovery. V12 (Carl Zeiss Microscopy, LLC, White Plains, NY, USA).

### Measurements of mosquito fecundity following aspirin treatment

To explore aspirin’s effect on mosquito oviposition and fecundity, individual female *An. gambiae* mosquitoes were transferred to 50 mL conical tubes containing a moistened filter paper at 48 h PBF. Oviposition rate was determined by observing the presence or absence of eggs every 24 h from 72 h PBF to 120 h PBF. At 120 h PBF, egg papers were fully submerged in water and allowed to hatch overnight. At 144 h PBF, total number of laid eggs and larval hatching rate were assessed.

### Ovarian developmental analysis using fluorescence microscopy

Ovaries from females (blood-fed) were dissected in 1xPBS and fixed with 3.7% paraformaldehyde in a wet chamber under darkness at room temperature (RT) for 60 min. After washing three times with 1xPBS, cells in ovarioles were permeabilized with 0.2% Triton X-100 in 1xPBS at RT for 20 min. Cells were then washed three times with 1xPBS and blocked with 5% skim milk (MB cell, Seoul, Korea) in 1xPBS at RT for 60 min. After washing once with 1xPBS, ovarian cells were incubated with FITC-tagged phalloidin in 1xPBS at RT for 1 h. After washing three times with 1xPBS, cells were incubated with DAPI (1 mg/mL) diluted 1,000 times in PBS at RT for 2 min for nucleus staining. After washing three times with 1xPBS, ovarian cells were observed under a fluorescence microscope (DM2500, Leica, Wetzlar, Germany) at 200× magnification.

### Bioinformatics analysis of PGE_2_ receptor (Aa-PGE_2_R) of *Ae. Albopictus*

DNA sequences of *Aa-PGE*_*2*_*R* were obtained from GenBank (www.ncbi.nlm.nih.gov) with an accession number of XM_029880552.1. Its translated protein sequence was obtained with an ExPASy translation tool (http://web.expasy.org/translate/). The N-terminal signal peptide was determined using SignalP 4.0 server (http://www.cbs.dtu.dk/services/SignalP/). Protein domain was predicted using Prosite (http://prosite.expasy.org/), Pfam (http://pfam.xfam.org), and InterPro (http://www.ebi.ac.uk/interpro/). Phylogenetic tree was generated with a Neighbor-Joining method using MEGA 6.

### RNA extraction and cDNA preparation

Total RNAs were extracted from different developmental stages (∼150 embryo, 50 young larvae and one adult both from male and female) of mosquitoes for developmental expression analysis. To analyze adult body parts, females at 8 days PE were used to isolate head, thorax, ovary, and abdomen. To collect total RNAs from blood-fed ovaries, 5 days old females were allowed to feed blood. From these samples, total RNAs were extracted using Trizol reagent (Invitrogen, Carlsbad, CA, USA) according to the manufacturer’s instruction. After RNA extraction, RNA was resuspended in nuclease-free water and quantified using a spectrophotometer (NanoDrop, Thermo Fisher Scientific, Wilmington, DE, USA). RNA (1 µg) was used for cDNA synthesis with RT PreMix (Intron Biotechnology, Seoul, Korea) containing oligo dT primer according to the manufacturer’s instructions.

### RT-PCR and RT-qPCR

RT-PCR was performed using Taq DNA polymerase (GeneALL, Seoul, Korea) with the following conditions: initial denaturation at 94°C for 5 min, followed by 35 cycles of 95°C denaturation for 30 sec, 57°C for 30 sec, 72°C extension for 30 sec, and a final extension at 72°C for 10 min. Each RT-PCR reaction mixture (25 µL) consisted of cDNA template, dNTP (each 2.5 mM), 10 pmol for each forward and reverse primer (**Table S1**), and Taq DNA polymerase (2.5 units/µL). Expression of actin (DQ657949.1) was used as reference.

All gene expression levels in this study were determined using a Real time PCR machine (Step One Plus Real-Time PCR System, Applied Biosystem, Singapore) under the guideline of Bustin et al. [22]. Real time PCR was conducted with a reaction volume of 20 µL containing 10 µL of Power SYBR Green PCR Master Mix (Thermo Scientific Korea), 5 µL of cDNA template (50 ng), and each 1 µL of forward and reverse primers (**Table S1**). After an initial heat treatment at 95°C for 10 min, qPCR was performed with 40 cycles of denaturation at 95°C for 30 sec, annealing at 57°C for 30 sec, and extension at 72°C for 30 sec. Expression level of actin was used as reference to normalize target gene expression levels under different treatments. Quantitative analysis was performed using the comparative CT (2^-ΔΔCT^) method [23].

### RNA interference (RNAi) against Aa-PGE_2_R

Template DNA was amplified with gene-specific primers containing T7 promoter sequence (5’-TAATACGACTCACTATAGGGAGA-3’) at the 5’ end. The resulting PCR product was used to *in vitro* synthesize double-stranded RNA (dsRNA) encoding *Aa-PGE*_*2*_*R* using T7 RNA polymerase with deoxynucleotide triphosphate mixture at 37°C for 3 h. dsRNA was mixed with a transfection reagent Metafectene PRO (Biontex, Plannegg, Germany) at 1:1 (v/v) ratio and incubated at 25°C for 30 min to form liposomes.

In injection experiment, 1 µg of dsRNA was injected into 5 days old females (10 min before BF) using PV830 microinjector under SZX-ILLK200 stereomicroscope. A control dsRNA (‘dsCON’) was prepared with the method of Vatanparast et al. [24] using a Megascript RNAi kit (Ambion, Austin, TX, USA). To prepare dsCON, a 500 bp fragment of *green fluorescence protein* (*GFP*) gene was synthesized. After quantification using Nanodrop Lite spectrophotometer, dsRNA was mixed with transfection reagent Metafectene PRO (Biontex, Plannegg, Germany) at 1:1 (v/v) ratio. For each treatment, three females were mated with one male.

### Analysis of RNAi-treated females and PGE_2_ rescue experiment

For RNAi treatment, 1 μg of gene-specific dsRNA in 1 μL was injected to 5 days PE females at 10 min before BF. After BF, females were reared with 10% sugar solution. To rescue RNAi-treated females, 1 μL of PGE_2_ (10 μg) was injected to females at 12 h after dsRNA injection. Control RNAi (‘dsCON’) was injected at the same concentration. Females at 72 h after BF were used to count the total number of oocytes under a stereo microscope (Stemi SV11, Zeiss, Germany). For fecundity test, three females were mated with one male in a replication. Each treatment was replicated three times.

### Observation of egg shell using scanning electron microscopy (SEM)

Eggs of *Ae. albopictus* were obtained from ASP-fed females at 48-72 h after oviposition and fixed in a mixture of 4% paraformaldehyde and 0.1% glutaraldehyde in 0.1 M sodium cacodylate buffer (pH 7.4) for 24 h at room temperature. Samples were then washed three times for 20 min with 0.1 M sodium cacodylate buffer and followed by dehydrated in a progressive ethanol gradient of 50, 60, 70, 90 and 100% for 20 min each. After drying, eggs were observed under scanning electron microscope (JSM-6300, Jeol, Tokyo, Japan).

### Gene expression analysis of choriogenesis-associated genes: *Yellow-g* and *Yellow-g2*

To analyze the molecular influence of ASP on chorion formation, two exochorion-darkening genes (*Yellow-g* and *Yellow-g2*) of *A. albopictus* were assessed by RT-qPCR as described above. Total RNAs were isolated from females (one female per replication) provided with 10% sucrose solution containing ASP (1 µg/µL) at 48 h BF. Real time PCR was conducted as mentioned previously. After an initial heat treatment at 95°C for 10 min, qPCR was performed with 40 cycles of denaturation at 95°C for 30 sec, annealing at 60°C for 30 sec, and extension at 72°C for 30 sec. Primers used the sequences described in Noh et al. [25]. Each treatment was replicated three times.

### Statistical analysis

All results are expressed as mean ± standard deviation. They were plotted using Sigma plot (Systat Software, San Jose, CA, USA). Means were compared by least square difference (LSD) test of one-way analysis of variance (ANOVA) using PROC GLM of SAS program [26] and discriminated at Type I error = 0.05.

## Results

### Egg development of *Ae. albopictus* females after BF

Newly emerged females of *Ae. albopictus* have previtellogenic oocytes (**Fig 1A**), where follicles develop after adult emergence. Each follicle contains several nuclei, and increase in size with time after adult emergence. Vitellogenic oocytes were clearly visible after BF along with an increase in ovary size. Each follicle is surrounded by the follicular epithelium and contains an oocyte and several nurse cells. Follicles grew slightly after adult emergence and rapidly increased in size soon after BF (**Fig 1B**). At 24 h post feeding, oocytes began to be enclosed with a chorion (**Fig 1C**). At 3 days post feeding, all oocytes were fully developed into chorionated oocytes and ready for oviposition.

**Fig. 1.**
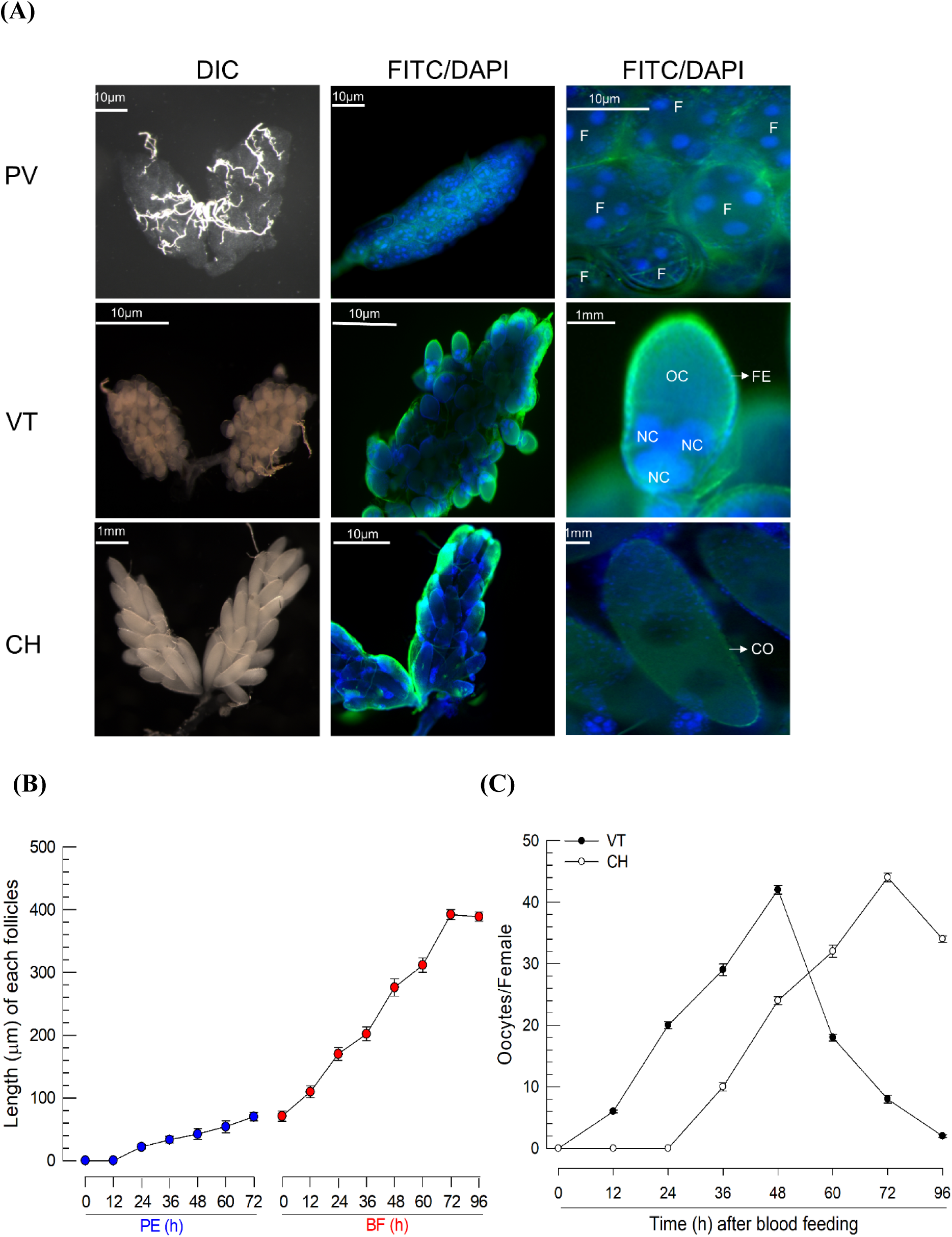
Oocyte development of *Ae. albopictus* females. (A) Three developmental stages of oocytes: previtellogenic (‘PV’), vitellogenic (‘VT’), and chorionated (‘CH’). Ovaries were observed at differential interference contrast (‘DIC’). Their cells were stained with FITC-tagged phalloidin (green) and DAPI (blue). Follicles (‘F’) were visible at PV stage. At VT stage, oocyte (‘OC’) and nurse cell (‘NC’) are enclosed with follicular epithelium (‘FE’). At CH stage, oocyte is enclosed with chorion (‘CO’). (B) Follicle growth in ovaries post-adult emergence (‘PE’) or post-blood feeding (‘BF’). (C) Oocyte development. Total ‘VT’ and ‘CH’ oocytes were calculated under a stereomicroscope at selected time points up to 96 h post-BF. For each treatment, 10 females were used in follicle growth and oocyte development.

### Aspirin injection inhibits choriogenesis

To determine the role of PG in mosquito choriogenesis, different eicosanoid biosynthesis inhibitors were injected to females just before BF (**Fig 2**). Dexamethasone, a general inhibitor of eicosanoid biosynthesis, significantly (*P* < 0.05) impaired the production of chorionated eggs. Two PG biosynthesis inhibitors, aspirin (ASP) and ibuprofen, also significantly (*P* < 0.05) inhibited the choriogenesis, whereas naproxen, an LT biosynthesis inhibitor, did not (**Fig 2A**). Addition of PGE_2_ rescued the choriogenesis of females treated with ASP (**Fig 2B**). The inhibitory activity of ASP on choriogenesis resulted in a significant (*P* < 0.05) reduction in the number of oviposited eggs (**Fig 2C**). The reduced fecundity was rescued by the addition of PGE_2_, but not by other types of eicosanoids such as 14,15-EET and LTB_4_. The oviposited eggs from the ASP-treated females also suffered poor hatch rates (**Fig 2D**). The reduced egg hatch rate was significantly (*P* < 0.05) rescued by the addition of PGE_2_.

**Fig. 2.**
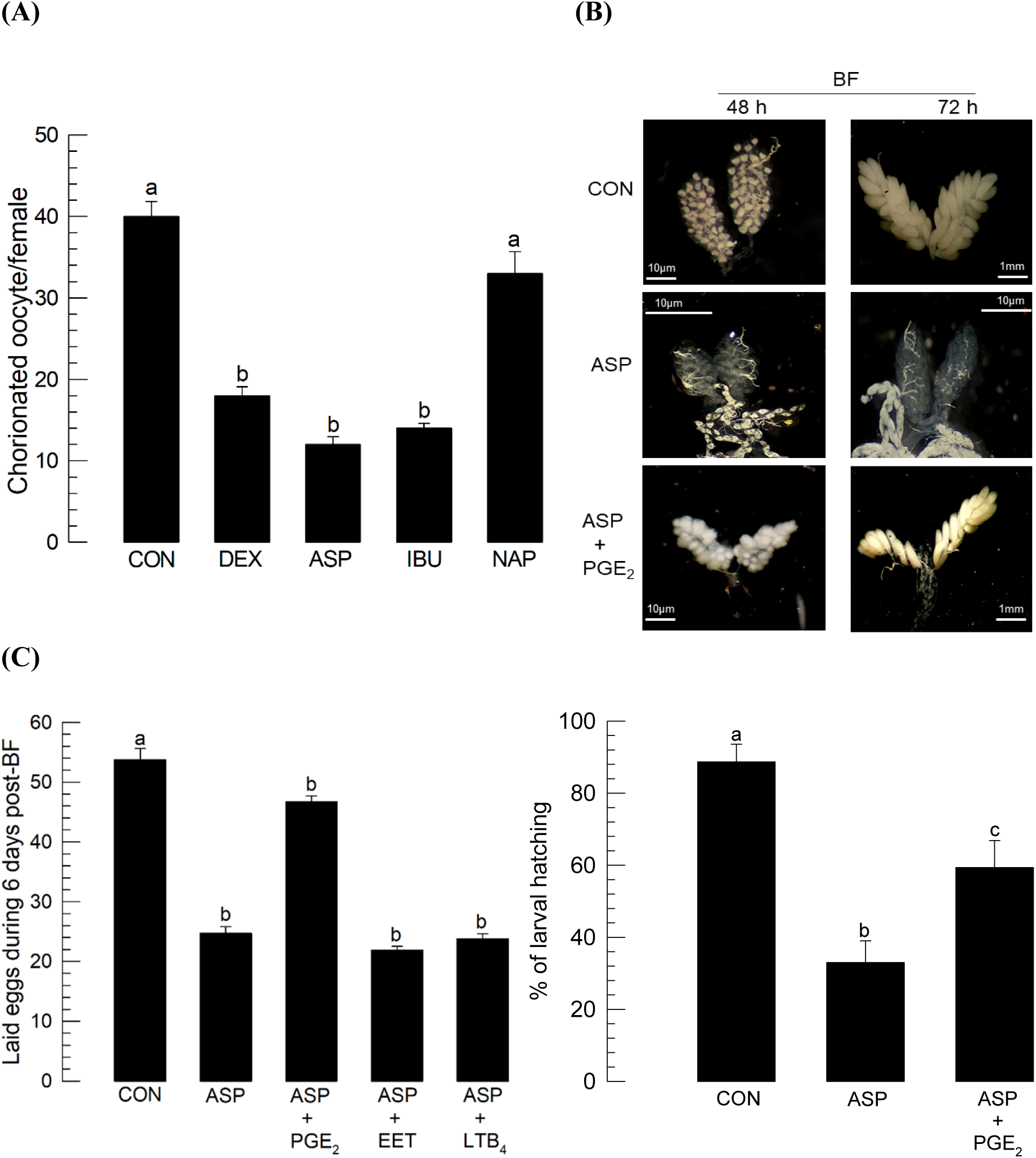
Effects of different eicosanoid biosynthesis inhibitors on egg production of *Ae. albopictus*. PLA_2_ inhibitor (dexamethasone: DEX), two COX inhibitors (ibuprofen: IBU, aspirin: ASP), and LOX inhibitor (naproxen: NAP) were used to treat 5 days old females before blood-feeding (‘BF’). At 10 min before BF, the inhibitor was injected to females at a dose of 1 μg/individual. Dimethylsulfoxide used to dissolve inhibitors was injected as a control (‘CON’). (A) Inhibitor effect on the number of chorionated oocytes at 72 h post BF. Each treatment was assessed using 10 females. (B) Rescue effect of PGE_2_ (10 μg/individual) on choriogenesis of females treated with ASP (1 μg/individual). Ovaries were observed at 48 and 72 h post-BF. (C) Specific rescue effect of PGE_2_ on choriogenesis of females treated with ASP. ASP was injected at a dose of 1 μg/individual. PGE_2_, 14,15-EET, or LTB_4_ was injected at 10 μg/individual to females treated with ASP. After BF, three females were mated with one male in each replication. Each treatment was replicated three times. The number of laid eggs was counted for 6 days after BF. (D) Decrease in hatch rate of the eggs laid by the ASP (1 µg/individual)- treated females and rescue effect of the addition of PGE_2_ (10 µg/individual). Each replication used 100 laid eggs. Each treatment was replicated three times. Different letters above standard deviation bars indicate significant difference among means at Type error = 0.05 (LSD test).

Inhibitory activities of ASP at different concentrations against choriogenesis were assessed, with results shown in **Fig. 3**. As expected, ASP inhibited choriogenesis of *Ae. albopictus* in a dose-dependent manner. When ASP was injected to females, only 10 ng of ASP per female was required to significantly (*P* < 0.05) reduce choriogenesis (**Fig 3A**). At 1 μg of ASP, about 80% oocytes failed to be chorionated compared to control. Oral administration of ASP also inhibited choriogenesis in a dose-dependent manner, although its inhibitory effect was much less than that of the injection method (**Fig 3B**). Even at 1,000 ppm of ASP, only 30% oocytes did not form chorions.

**Fig. 3.**
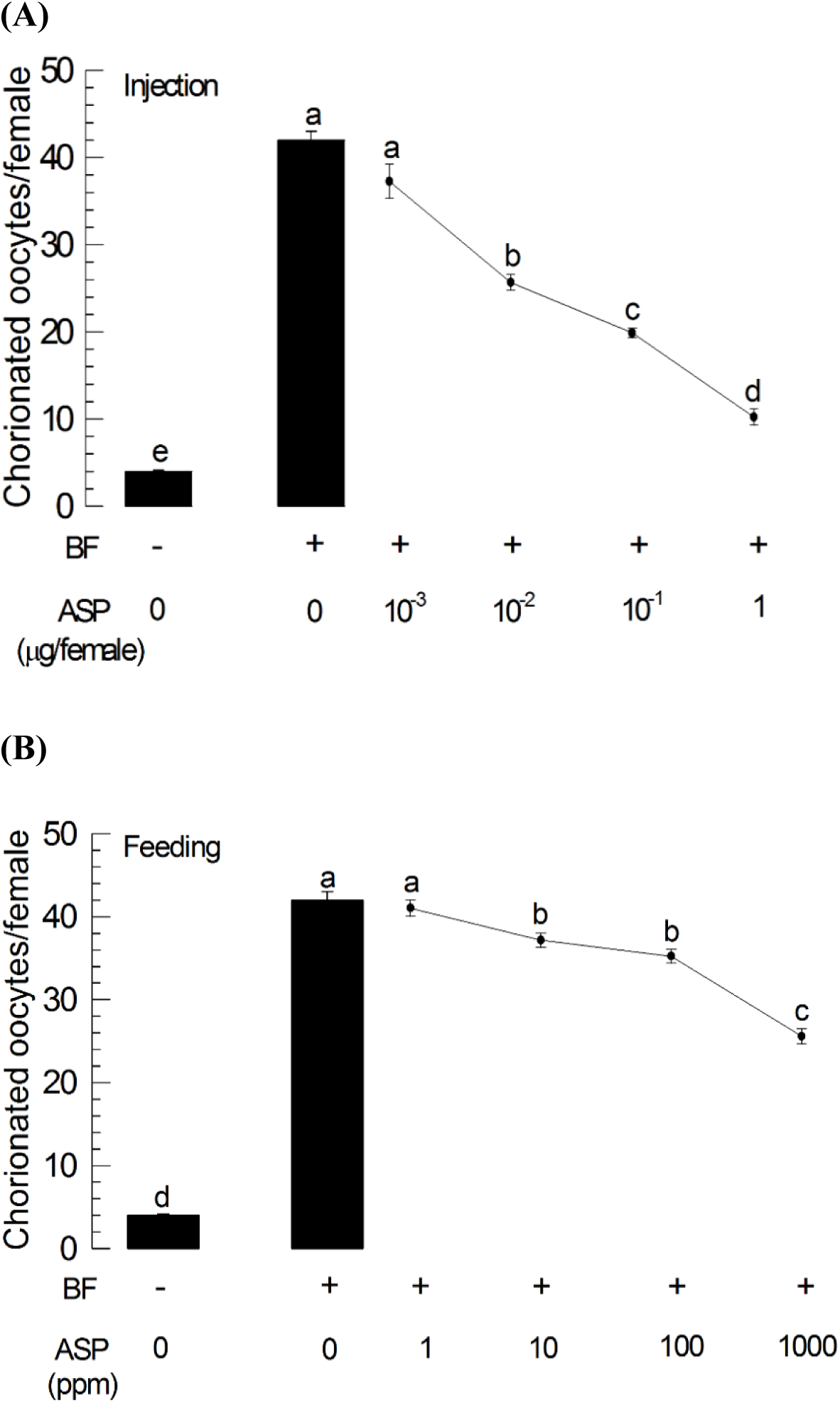
Effects of different concentrations of aspirin (‘ASP’) on choriogenesis of *Ae. albopictus*. (A) Hemocoelic injection. ASP was injected to 5 days old females at 10 min before blood feeding (‘BF’). At 72 h post BF, ovaries were collected from treated females in PBS to count the number of chorionated oocytes. (B) Oral feeding. Different concentrations of ASP were prepared with 10% sugar solution. The sugar-ASP solution (5 mL) was provided to an adult cage containing 10 females at 3 h before BF. After BF, females were allowed to continuously feed the sugar-ASP solution for 3 days and then assessed for the development of chorionated oocytes. Each treatment used 10 females. Different letters above standard deviation bars indicate significant difference among means at Type error = 0.05 (LSD test).

### Aspirin effect on egg production of *An. gambiae*

Similar to the detrimental effects on reproduction and larval hatching by oral administration of ASP in *Ae. albopictus*, we observed that the administration of ASP at 10 mg/mL by oral feeding in *An. gambiae* also resulted in the attenuated choriogenesis of the ovarian follicle at 72 h PBF (**Fig 4A**). ASP feeding significantly reduced the number of chorionated oocytes in *An. gambiae* as compared to sucrose control (**Fig 4B**). Moreover, ASP negatively influenced mosquito fecundity, causing a reduction in the number of oviposited eggs (**Fig 4C**) as well as a decrease in larval hatching (**Fig 4D**) at 144 h PBF.

**Fig. 4.**
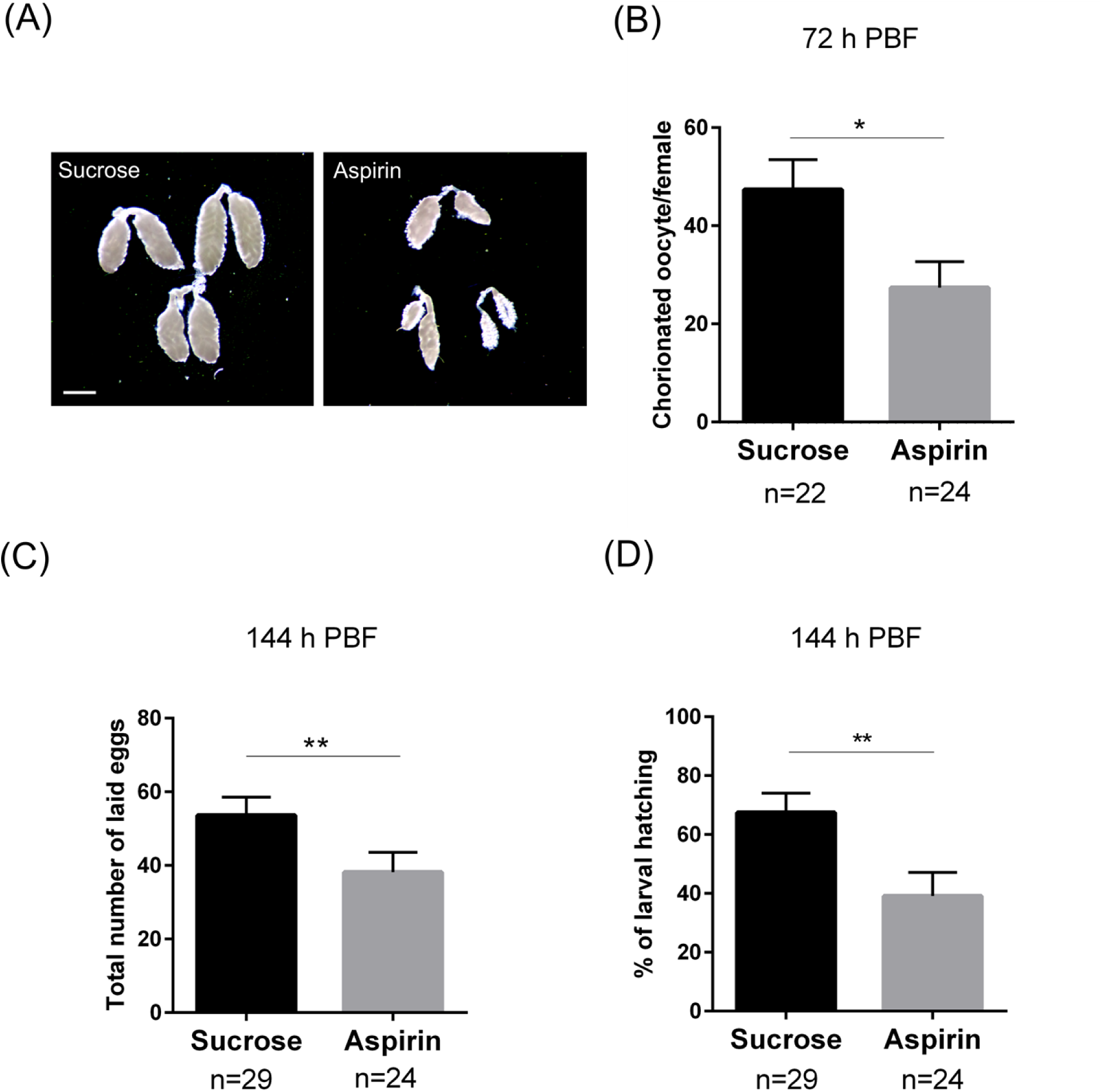
Effects of aspirin feeding on *An. gambiae* choriogenesis, fecundity and larval hatching. (A) Dissected ovaries from mosquitoes fed either aspirin (10 mg/mL) or 10% sucrose solution at 72 h post-blood feeding (PBF). Scale bar represents 1 mm. (B) Aspirin feeding significantly limited development of chorionated oocytes as compared to 10% sucrose control. (C) Aspirin feeding impaired mosquito fecundity, resulted in less number of laid eggs at 144 h PBF. (D) A significant decrease of larval hatching rate was observed with aspirin feeding at 144 h PBF. Data were analyzed by a Mann–Whitney test using GraphPad Prism 6.0. Bars represent mean ± SE of three independent replications. Asterisks denote significance (\**P* < 0.05, \*\**P* < 0.01).

### Aspirin suppresses Yellow genes associated with chorion formation

To understand a molecular action of ASP to inhibit choriogenesis of mosquitoes, expression of *Yellow-g* and *Yellow-g2*, which are required for exochorion darkening and subsequent rigidity in *Ae. albopictus* [25], was assessed after inhibitor treatment (**Fig 5**). Most eggs collected from ASP-treated mosquitoes exhibited a malformed egg shell as observed from SEM pictures (**Fig 5A**). ASP treatment significantly (*P* < 0.05) suppressed expression of both Yellow genes (**Fig 5B**).

**Fig. 5.**
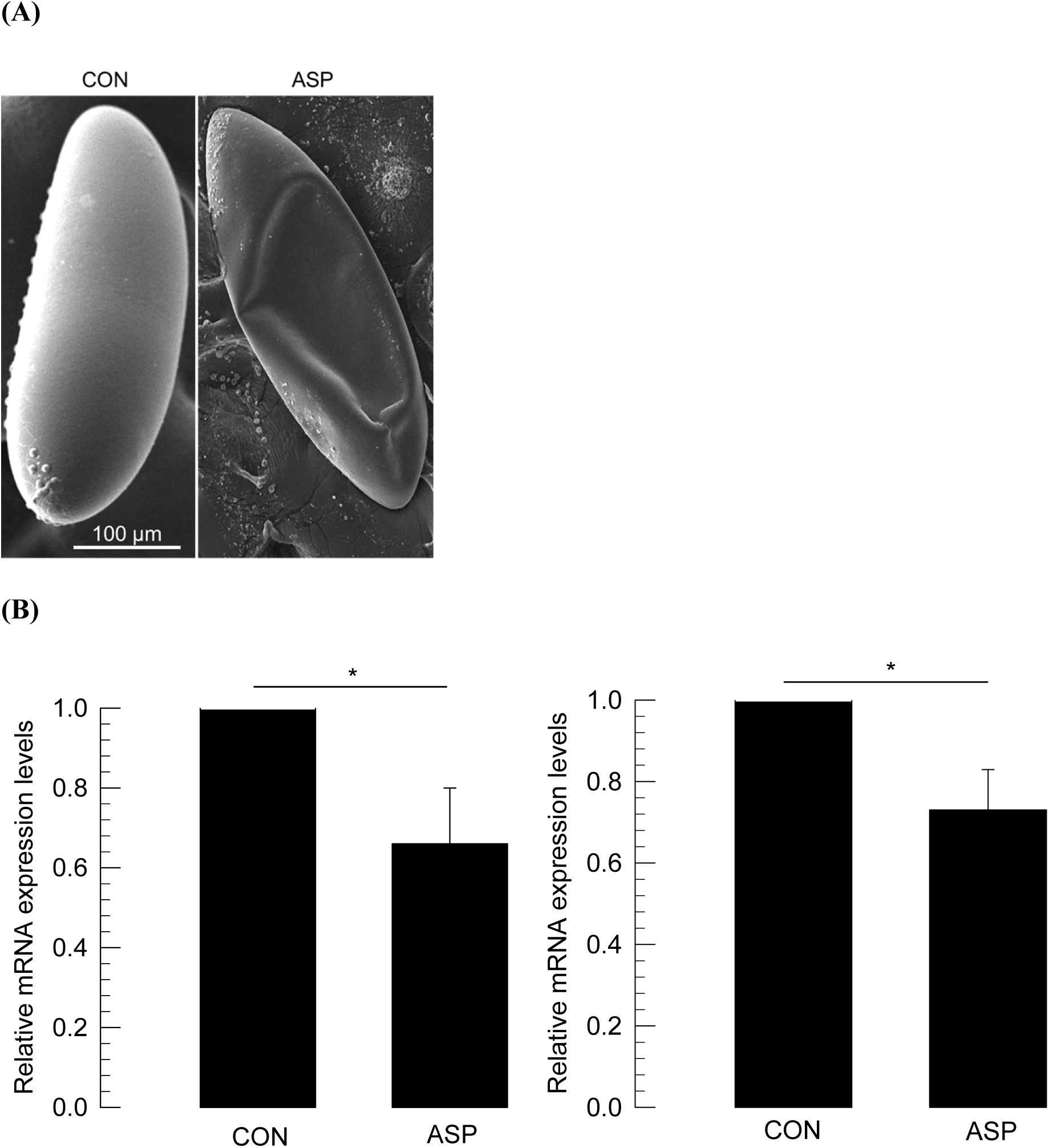
Inhibitory effect of aspirin (‘ASP’) on expression of Yellow genes (*Yellow-g* and *Yellow-g2*) associated with exochorion formation. (A) SEM pictures demonstrating a malformed chorion of eggs laid by females treated with ASP (1 µg/µL) in 10% sucrose solution. Eggs were collected at 48-72 h after oviposition and used for SEM imaging. (B) RT-qPCR of *Yellow-g* (left panel) and *Yellow-g2* (right panel). All treatments were independently replicated three times. Asterisk mark above standard deviation bars indicates significant difference between means at Type I error = 0.05 (LSD test).

### Identification of PGE_2_ receptor of *Ae. albopictus*

The stimulating effect of PGE_2_ on choriogenesis suggests that there is a PGE_2_ receptor on follicles of *Ae. albopictus*. Interrogation of *Ae. albopictus* with the *M. sexta* PGE_2_ receptor sequence [27] allowed us to predict a PGE_2_ receptor gene (*Aa-PGE*_*2*_*R*) in *Ae. albopictus* (**Fig 6**). *Aa-PGE*_*2*_*R* encoded 398 amino acid residues and seven transmembrane domains (**Fig 6A**). Its predicted amino acid sequence was clustered with other EP4 receptors (**Fig 6B**). *Aa-PGE*_*2*_*R* was expressed in all developmental stages. It was highly expressed in adult females (**Fig 7A**). In female adults, it was highly expressed in ovary tissue (**Fig 7B**). Expression of *Aa-PGE*_*2*_*R* in ovaries increased along with increasing number (r = 0.9972; *P* < 0.0001) of chorionated oocytes (**Fig 7C**).

**Fig. 6.**
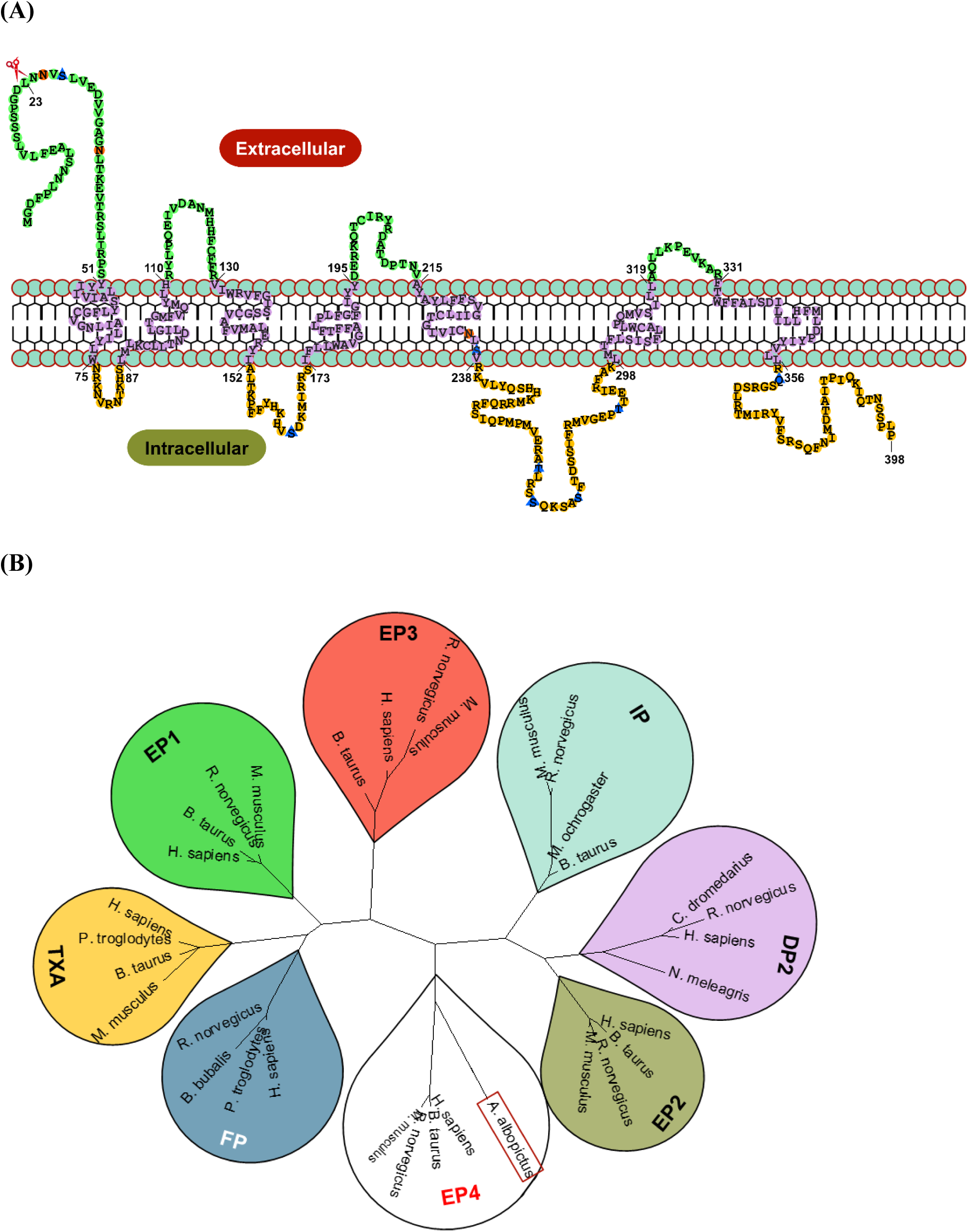
Molecular characteristics of *Ae. albopictus* PGE_2_ receptor (*Aa-PGE*_*2*_*R*: GenBank accession number: XP_029736412.1). (A) Domain analysis. A cleavage site of signal peptide (23 residues) is denoted by scissor. Seven transmembrane domains are marked on phospholipid bilayer with locations. N-glycosylation sites are denoted by orange circle and phosphorylation sites are marked by blue triangle. (B) A phylogenetic analysis. The tree was generated by the Maximum likelihood tree method using software package MEGA6.0, where evolutionary distances were computed using Poisson correction method. Amino acid sequences of different PGE_2_R were retrieved from GenBank with accession numbers of KAB1277535.1 for *C. dromedarius* (DP2), NP_000944.1 for *H. sapiens* (DP2), NP_071577.1 for *R. norvegicus* (DP2), XP_021258611.1 for *N. meleagris* (DP2), AAI12966.1 for *H. sapiens* (FP), JAA36212.1 for *P. troglodytes* (FP), AAZ08074.1 for *B. bubalis* (FP), EDL82480.1 for *R. norvegicus* (FP), NP_032993.2 for *M. musculus* (IP), NP_001015622.2 for *B. Taurus* (IP), EDM08294.1 for *R. norvegicus* (IP), XP_005361096.1 for *M. ochrogaster* (IP), NP_001345441.1 for *M. musculus* (TXA), NP_001161391.1 for *B. Taurus* (TXA), JAA33130.1 for *P. troglodytes* (TXA), AAH74749.1 for *H. sapiens* (TXA), NP_000946.2 for *H. sapiens* (EP1), NP_001265404.1 for *R. norvegicus* (EP1), NP_038669.1 for *M. musculus* (EP1), NP_001179077.1 for *B. taurus* (EP1), NP_000947.2 for H. sapiens (EP2), NP_112350.1 for *R. norvegicus* (EP2), NP_032990.1 for *M. musculus* (EP2), NP_777013.1 for *B. taurus* (EP2), NP_942007.1 for *H. sapiens* (EP3), NP_036836.1 for *R. norvegicus* (EP3), NP_001346674.1 for *M. musculus* (EP3), NP_851375.1 for *B. taurus* (EP3), NP_000949.1 for *H. sapiens* (EP4), NP_114465.3 for *R. norvegicus* (EP4), NP_001129551.1 for *M. musculus* (EP4), and NP_777014.1 for *B. Taurus* (EP4).

**Fig. 7.**
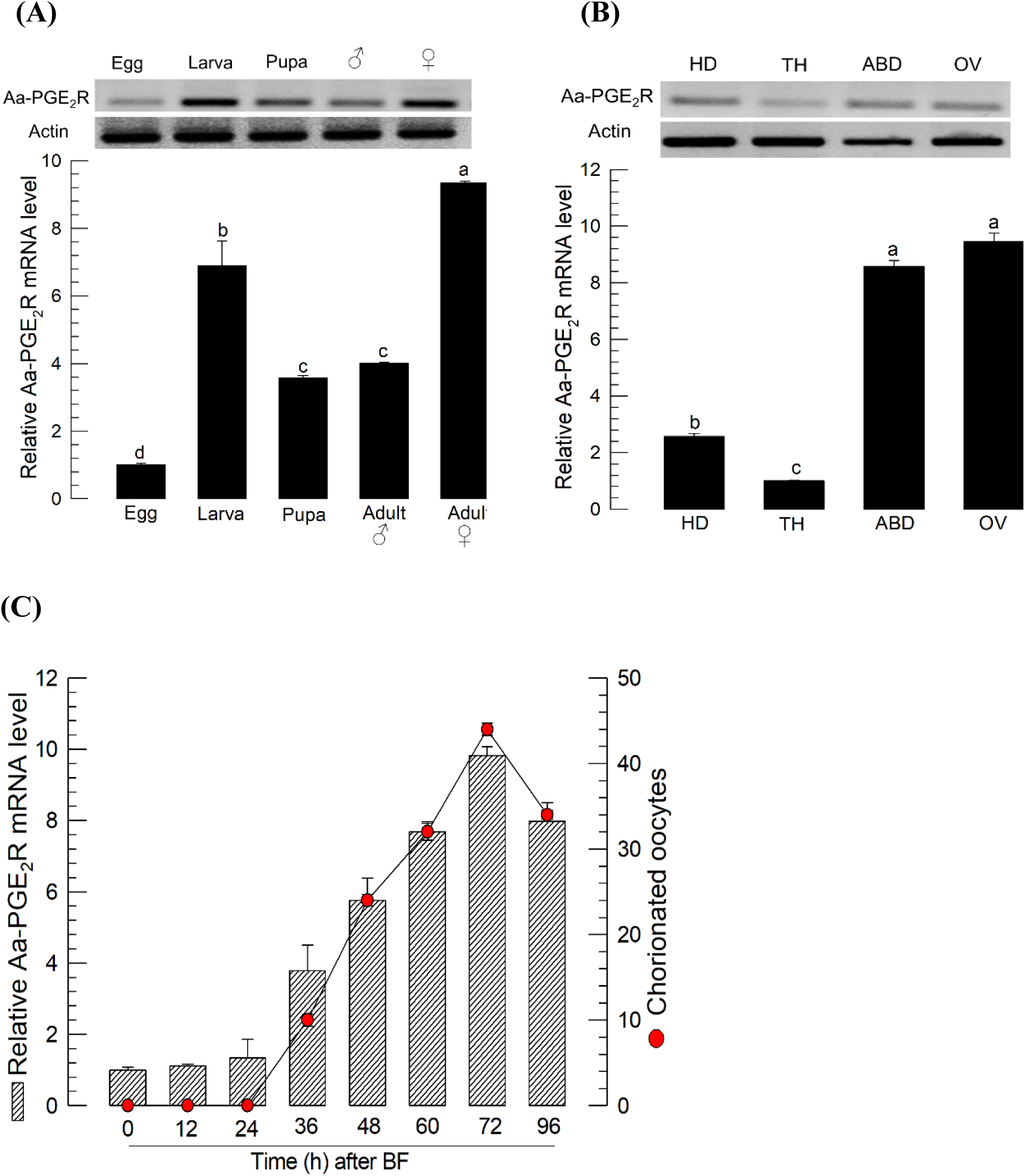
Expression profile of *Aa-PGE*_*2*_*R*. (A) Expression in different developmental stages from egg to male and female adults. (B) Expression in different body parts of *Ae. albopictus* females at 3 days after blood-feeding (‘BF’): head (‘HD’), thorax (‘TH’), ovary (‘OV’), and abdomen (‘ABD’). (C) Relationship between the expression level of *Aa-PGE*_*2*_*R* and the number of chorionated oocytes after BF. Expression of actin, a reference gene, was used to normalize gene expression level in RT-qPCR. All treatments were independently replicated three times. Different letters above standard deviation bars indicate significant difference among means at Type I error = 0.05 (LSD test).

### RNA interference of *Aa-PGE*_*2*_*R* expression inhibits choriogenesis

Double-stranded RNA specific to *Aa-PGE*_*2*_*R* significantly (*P* < 0.05) suppressed its expression level when it was injected to teneral female adults (**Fig 8A**). RNAi-treated adults failed to develop chorionated oocytes as compared to control females (**Fig 8B**). PGE_2_ addition to RNAi-treated females did not alter the reduction in choriogenesis.

**Fig. 8.**
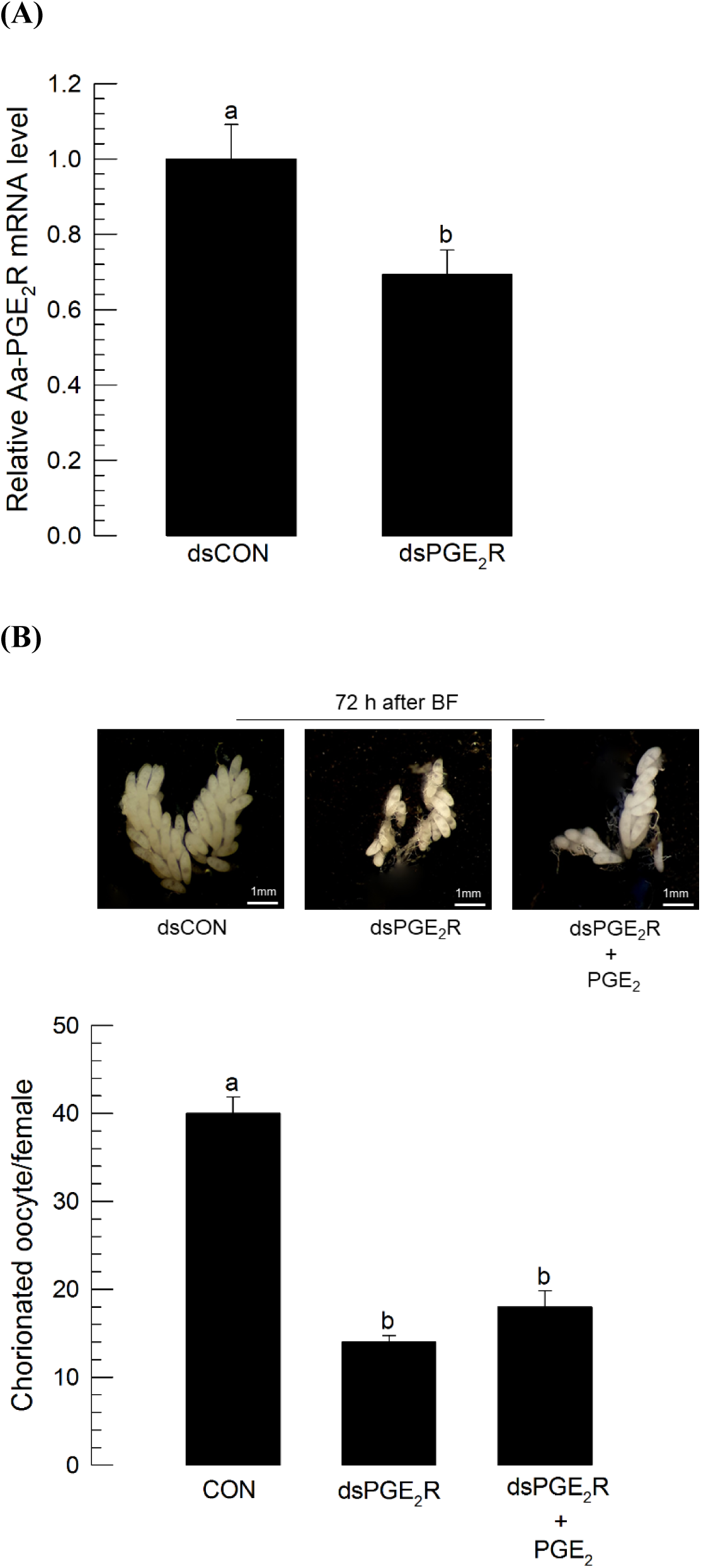
Effect of RNA interference (RNAi) of *Aa-PGE*_*2*_*R* on choriogenesis of *Ae. albopictus*. RNAi of *Aa-PGE*_*2*_*R*. RNAi was performed by injecting 1 μg of gene-specific dsRNA (‘dsPGE_2_R’) or control dsRNA (‘dsCON’) to 5 days old females at 10 min before blood-feeding (BF). Changes in mRNA levels were assessed at 72 h after injecting dsPGE_2_R or dsCON by RT-qPCR using actin as a reference gene to normalize target gene expression level. Each treatment was replicated three times. (B) Influence of RNAi on choriogenesis. At 12 h after dsRNA injection, 1 µL of PGE_2_ (10 μg per female) was injected to RNAi-treated adults. At 72 h after BF, total number of chorionated oocytes was counted under a stereomicroscope. Each treatment used 10 females. Different letters above standard deviation bars indicate significant difference among means at Type I error = 0.05 (LSD test).

## Discussion

In insects, the first report on the physiological role of PG was regarding the egg laying behavior of house cricket (*Acheta domesticus*) females [28]. In the present study, aspirin-fed females during the last nymphal stage were impaired in fecundity, suggesting a role of PG in female reproduction. Loher et al. [29] analyzed PGE_2_ levels in females and found a significant increase of hormone levels after mating. The PGE_2_ level in the mated females are much higher than the level in the male reproductive organ. This suggested a hypothesis of PG-synthetic machinery transfer during mating, in which males transfer PGE_2_-biosynthetic enzymes in the spermatophores and subsequently the mated females synthesize PGE_2_ in the spermathecae to release PGE_2_ to hemolymph. In addition, Loher et al. [8] showed that cricket spermatophores had PGE_2_ biosynthetic activity which was specifically inhibited by aspirin. For PGE_2_ synthesis from arachidonic acid as a precursor, COX activity is necessary to form PGH_2_ via *bis*-dioxygenation and subsequent reduction [30]. However, no COX ortholog has been identified in insect genomes [31]. Instead, insects are likely to use COX-like peroxinectin (Pxt) [15]. Two Pxts have been identified in *S. exigua*. They can mediate PG biosynthesis [16]. For PGE_2_ synthesis (PGES), an isomerase activity to change PGH_2_ to PGE_2_ is required in *S. exigua* [32]. Based on identification of similar Pxts in another mosquito, *An. gambiae* [17], *Ae. albopictus* might have a similar ability to biosynthesize PGE_2_. Indeed, PGE_2_ has been detected in other mosquitoes. For example, specific stellate cells of Malpighian tubules possess PGE_2_ in *Ae. aegypti* to mediate fluid secretion [33]. PGE_2_ production identified in the hemolymph of *An. gambiae* was significantly increased following *Plasmodium* infection [17]. In *An. albimanus*, PGE_2_ has been detected in the midgut. It plays a crucial role in modulating gene expression of antimicrobial peptides [20]. Based on these findings of PGE_2_ in mosquitoes, this study tested the role of PG in mediating choriogenesis of *Ae. albopictus* and *An. gambaie*. Treatment with PG biosynthesis inhibitor and interference of PGE_2_ signal using RNAi bolstered the conserved role of PGs in mediating egg development of mosquito choriogenesis.

Inhibition of PG biosynthesis using specific inhibitors suppressed the choriogenesis of *Ae. albopictus*. PGE_2_ addition significantly rescued the choriogenesis inhibited by aspirin treatment. PGs can function as autocrine mediators in mammals and insects in various physiological processes [13]. PGs play crucial roles in mediating oocyte development of *D. melanogaster* [11], *B. mori* [10], and *S. exigua* [18]. In both dipteran and lepidopteran insects, PGE_2_ or PGD_2_ can mediate choriogenesis by stimulating chorion-associated gene expression [10,19]. Machado et al. [10] have used an *in vitro* ovarian follicle culture of *B. mori* and shown that incubating follicular epithelium cells in the presence of PG biosynthesis inhibitors can block the transition from follicle development to choriogenesis. They have suggested that PGs can act on expression of chorion genes. Alternatively, PGs may play a role to changing egg development from vitellogenesis to choriogenesis. Thus, aspirin treatment might impair this transition, inhibiting choriogenesis in both *Ae. albopictus* and *An. gambiae*. In *Rhodnius prolixus*, PGs produced by the ovary can inhibit endocytosis of *Rhodnius* heme-binding protein by the oocyte, implying a functional role of PGs in the termination of vitellogenesis [34]. This might be explained by the volume increase of follicular epithelial cells to prevent its patency to uptake vitellogenin [35], which could be induced by cAMP that might be a secondary messenger of PGE_2_. Thus, PGs are likely to be required for choriogenesis in mosquitoes.

Two Yellow genes associated with exochorion formation were significantly suppressed in their expression by ASP treatment. Yellow and related major royal jelly protein (MRJP)-like proteins are widely found in insect genomes and these genes are classified into ten clades including Yellow-b, -c, -d/e3, -e, -f, -g/g2, -h, -y, -x and MRJP-like protein [36]. The two Yellow genes of *Ae. albopictus* are highly homologous molecular size (379 and 373 amino acid residues) except for four substitutions at 37th (D→H), 286th (L→M), 358th (C→W), and 364th (S→F) residues [37]. *Yellow-g* and *Yellow-g2* are specifically expressed in ovary after BF and their proteins are localized in exochorion and outer endochorion, at which they mediate darkening processes to make the chorions to be physically strengthened [25]. Thus, the suppression of these gene expressions by ASP treatment may lead to a malformed chorion as observed in this current study. This suggests that PGs modulate the expression of these two genes. However, the mechanism by which PGs influence gene expression needs to be investigated.

We identified the PGE_2_ receptor of *Ae. albopictus* and demonstrate its role in suppressing choriogenesis via RNAi, even with the addition of PGE_2_. Unlike the lepidopteran PGE_2_ receptors of *Manduca sexta* and *Spodoptera exigua*, which are classified into EP2 type [27,38], the predicted amino acid sequence of *Aa-PGE*_*2*_*R* shared homology with EP4 among four identified EP receptors. EP4 has been implicated in various physiological and pathological responses in mammals [39]. When bound to PGE_2_, EP4 can mobilize trimeric G proteins to be dissociated into Gαs and Gβγ components to stimulate adenyl cyclase to raise intracellular levels of cAMP. EP4 activation of G proteins can also activate PI3K/AKT/mTOR, ERK, and p38 MAPK pathways [40]. These findings suggest that Aa-PGE_2_R can modulate various physiological processes using these down-stream signal pathways in *Ae. albopictus*. For example, *Aa-PGE*_*2*_*R* was highly expressed in the larval and adult stage. This suggests that Aa-PGE_2_R can mediate physiological processes other than adult reproduction. Actin cytoskeletal rearrangement in hemocytes of *S. exigua* is mediated by PGE_2_ during hemocyte-spreading to perform cellular immune responses [18]. Actin filaments are rearranged in nurse cells in response to PGE_2_. They play a crucial role in nurse cell dumping during oogenesis of *D. melanogaster* [9]. During choriogenesis, a massive amount of protein synthesis occurs. Altering PG signaling using specific inhibitors can interrupt the expression of chorion-associated genes in *B. mori* [10]. The inhibition of choriogenesis by RNAi specific to *Aa-PGE*_*2*_*R* suggests that PG signaling is required for choriogenesis of *Ae. albopictus*, supporting the inhibitory activity of aspirin treatment observed with both *Ae. albopictus* and *An. gambiae*. The inhibitory effect of aspirin or RNAi on choriogenesis of *Ae. albopictus* might be induced by interfering with vitellogenesis. Mosquito vitellogenesis is mainly controlled by JH and 20E [41]. JH is usually released soon after adult emergence. It mediates the differentiation of fat body to be competent to synthesize vitellogenin in response to 20E by increasing ribosomal biogenesis [42]. After a blood meal, a female mosquito can synthesize Vg in response to 20E [43]. JH again activates follicular patency to facilitate Vg uptake by growing oocytes [7]. During oogenesis, however, nurse cells should be degenerated by secreting their contents to adjacent oocyte, also known as nurse cell dumping [44]. Such nurse cell dumping is mediated by PGs via cytoskeleton rearrangement by bundling actin filament through Fascin in *Drosophila* [9]. *Ae. albopictus* ovaries were observed to be composed of several polytrophic ovarioles in the current study. Thus, the nurse cell dumping might be required for oogenesis and subsequent vitellogenesis. Aspirin treatment or *Aa-PGE*_*2*_*R*-silencing can inhibit PG biosynthesis or block PG signaling in target cells, which might lead to attenuated egg development in mosquitoes.

The inhibition of egg development by aspirin also has great promise as a novel control strategy against important mosquito vector species. In the current study, orally administered aspirin interfered with egg development, demonstrating its potential application to reduce mosquito populations. Furthermore, PGs are likely to play crucial roles in gut immunity of mosquitoes against microbiota [17]. Inhibition of PG production by aspirin treatment might also impair mosquito immunity, making mosquitoes more susceptible to microbial pathogens. Thus, designing and synthesizing aspirin derivatives might have potential to develop highly potent mosquitocidal agents. In addition, adverse effects of aspirin on egg development and other physiological processes should be tested against other mosquitoes in the future.

## Acknowledgements

A laboratory colony of *Ae. albopictus* used in this study was kindly donated by Professor Donghun Kim (Kyungpook National University, Sangju, Korea).

## Author Contributions

**Conceptualization**: Yonggyun Kim

**Data curation**: Abdullah Al Baki, Shabbir Ahmed, Hyeogsun Kwon, David Hall

**Formal analysis**: Abdullah Al Baki, Shabbir Ahmed, Hyeogsun Kwon, David Hall

**Funding acquisition**: Ryan Smith, Yonggyun Kim

**Investigation**: Abdullah Al Baki, Shabbir Ahmed, Hyeogsun Kwon, David Hall

**Methodology**: Abdullah Al Baki, Shabbir Ahmed, Hyeogsun Kwon, David Hall

**Project administration**: Ryan Smith, Yonggyun Kim

**Resources**: Ryan Smith, Yonggyun Kim

**Software**: Abdullah Al Baki, Shabbir Ahmed, Hyeogsun Kwon, David Hall

**Writing – original draft**: Abdullah Al Baki

**Writing – review & editing**: Ryan Smith, Yonggyun Kim

## Supplementary data

**Table S1.**
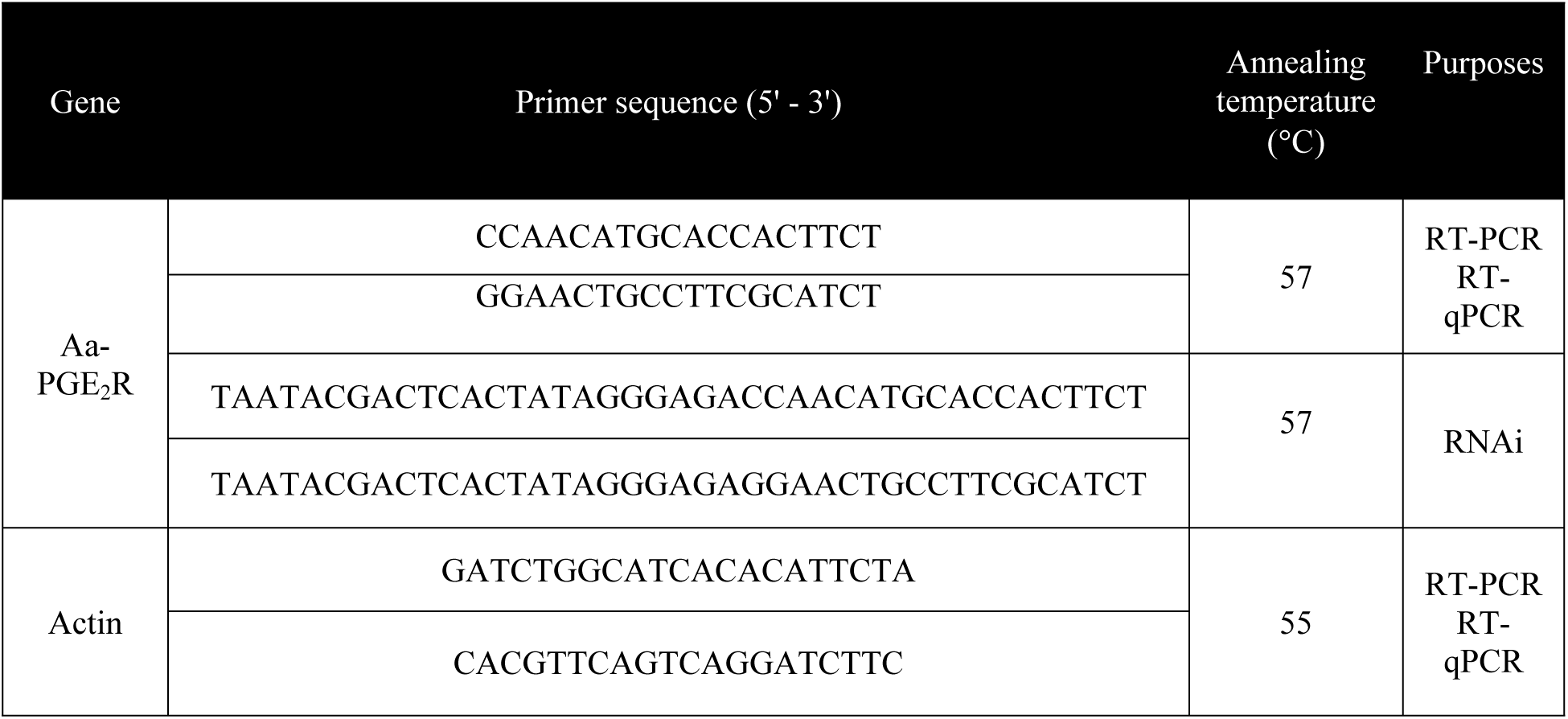
Primers used in this study

